# GEESE: Metabolically driven latent space learning for gene expression data

**DOI:** 10.1101/365643

**Authors:** Marco Barsacchi, Helena Andres Terre, Pietro Lió

## Abstract

Gene expression microarrays provide a characterisation of the transcriptional activity of a particular biological sample. Their high dimensionality hampers the process of pattern recognition and extraction. Several approaches have been proposed for gleaning information about the hidden structure of the data. Among these approaches, deep generative models provide a powerful way for approximating the manifold on which the data reside.

Here we develop GEESE, a deep learning based framework that provides novel insight into the manifold learning for gene expression data, employing a metabolic model to constrain the learned representation. We evaluated the proposed framework, showing its ability to capture biologically relevant features, and encoding that features in a much simpler latent space. We showed how using a metabolic model to drive the autoencoder learning process helps in achieving better generalisation to unseen data. GEESE provides a novel perspective on the problem of unsupervised learning for biological data.

**Availability:** Source code of GEESE is available at https://bitbucket.org/mbarsacchi/geese/.

## 1 Introduction

Metabolism comprises the complex network of chemical reactions allowing an organism to transform nutrients into energy and base components necessary for growth, replication, defence and other cellular tasks Metabolic reactions are mediated by enzymes which, in turn, are produced through gene expression. Nevertheless, despite this close association, the problem of predicting a metabolic phenotype from gene expression levels is anything but solved [29].

Gene expression microarrays provide a snapshot of all the transcriptional activity in a biological sample [27], which, in turn, is the result of the complex interactions among genes and environmental factors [12]. Gene expression profiling aims at capturing gene expression patterns in cellular responses to diseases, genetic perturbations and drug treatments. Nevertheless, while more than two decades have passed since the inception of the transcriptomic era, it is still easier to collect data that to understand it [18]. As a sample is typically characterised by large quantities of variables with unknown correlation structures, it is thus extremely challenging to analyse and make sense of the data. Dimensionality reduction techniques have been widely used to unravel the patterns hidden in the gene expression data [26]. As the difficulty of a learning task depends on how the information is represented, is of paramount importance to find a good representation for the data.

The *manifold hypothesis* states that high dimensional data tend to lie in the vicinity of a low dimensional manifold and represents the foundation of manifold learning [7]. Deep generative models, such as variational autoencoders (VAEs) [17] and generative adversarial networks (GANs) [9], involve learning a mapping (via a generator or decoder network) from a lower-dimensional latent space to the high-dimensional space of observed data [25]; in other words, they approximate the original manifold.

In this paper we propose an approach for learning structure in gene expression data; advancing the current state of the art, we employ the metabolic knowledge we possess in the form of a genome scale metabolic model (GSMM) [16], constraining the learnt representation to convey genotype-phenotype relationship. Furthermore, we endeavour to yield a disentangled representation, in which different latent units are sensible to changes in a single generative factor and approximately invariant with respect to changes of the other factors [2]. To do so, we employ a deep generative model (,0-VAE) [11] combined with a genome scale metabolic model. To the best of our knowledge, even if approaches aimed at extracting latent spaces from transcriptomic data [30] have been proposed before, none of them has integrated the metabolic model to constrain the learnt representation. We dubbed our framework GEESE (Gene Expression latEnt Space Encoder). The proposed approach could be somehow related to Relational Autoencoders [20], yet we are tackling related problems from a different and, in our case, more specific, perspective.

The rest of the work is organised as follows. In Section 2 we thoroughly describe the proposed framework as well as the theory and computational methods required; we then proceed to a detailed evaluation in Section 3. Conclusions are drawn in Section 4.

## 2 Methods

In this section we describe the proposed framework. Its schematic is provided in Figure 1. The general idea is introduced here; a detailed description is given in the following subsections.

**Figure 1.**
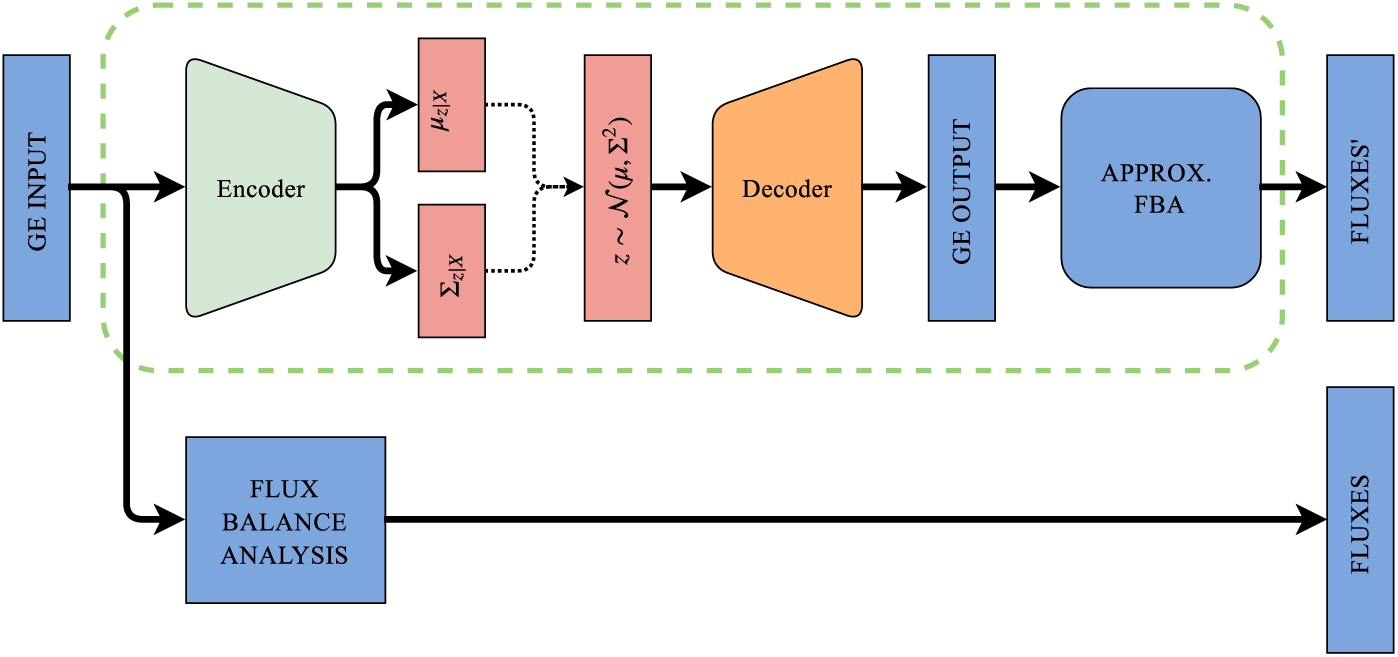
Global scheme of the proposed framework. The gene expression data is inputted both to the VAE and the FBA model. The VAE is trained minimizing the loss between the fluxes obtained by passing the reconstructed gene expression through the approximated FBA and the fluxes outputted by the real FBA.

Suppose we have a dataset 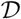 of gene expression data, from which we aim at extracting a (low dimensional) latent space representation. The dataset of gene expression data is provided as input to the FBA model, which produces a set of reaction fluxes *F*. We then train the FBA approximator, providing g.e. data as input and minimising the MSE between the fluxes *F* and the predicted fluxes *F*′. The VAE is then trained, with g.e. data as input, aiming at minimising the loss between the fluxes *F*′ obtained by passing the reconstructed gene expression through the approximated FBA and the fluxes *F*, keeping the weights of the FBA approximator fixed.

### 2.1 Genome Scale Metabolic Models

Metabolic models are based on a well-curated genome-scale metabolic network. A metabolic model comprises both the metabolic reactions and the genes encoding enzymes involved in mediating that reactions [19]. The metabolic network is topologically represented as **S**, a *M* × *N* stoichiometric matrix, where the *M* rows represent the stoichiometric coefficients of the corresponding metabolites in all the *N* reactions. Under the mass balance constraint, the system behaves according to a set of differential equations:

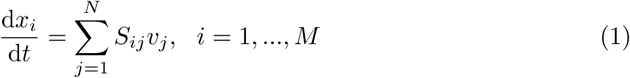

being *x_i_* the concentration of the *i*-th metabolite, and *v_j_* the flux through reaction *j* [24]. The value of *S_ij_* is negative, positive or zero if the *i*-th metabolite acts as reactant, product or does not participate in the *j*-th reaction respectively. Under the hypothesis of pseudo steady-state 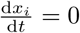, ∀*i*, the amount of compound being produced equals the total amount being consumed, thus:

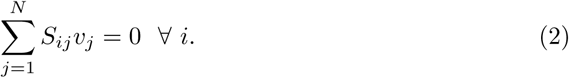

The pseudo steady-state hypothesis holds under the assumption that the time constants for metabolic reactions (milliseconds to tens of seconds) are typically much smaller than those of other cellular processes such as transcriptional regulation (minutes) or cellular growth (several minutes or hours) [28].

As there are more unknown variables than equations (*N ≥ M*), the system is undetermined and there are an infinite number of flux vectors that satisfy Eq. 2; that set of vectors represent the *null space* of **S**. In order to constrain the set of allowable metabolic phenotypes of the systems, thermodynamic and capacity constraints can be introduced into the system, as lower and upper bounds on the reaction fluxes: **v***^inf^* ≤ **v** ≤ **v***^sup^*. The mass-balance constraints, along with the flux inequalities define a convex-bounded polytope, in which all feasible solutions lie. The maximisation of a postulate objective function can be used to select the most physiologically relevant metabolic phenotype, leading to the classic FBA optimisation formulation [15]:

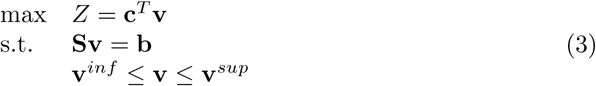

The most widely used objective function is the biomass function; it is a fictitious reaction used as a sink for biomass precursors (e.g. DNA, RNA, proteins, lipids) that together define the biomass composition of the cell [23]. The biomass objective relies on the assumption that the cell is striving to maximise its growth, given a fixed amount of resources.

Nonetheless, the usage biomass as the objective imposes a strong limitation, as an organism is often simultaneously optimising multiple and competing objectives [22]. Furthermore, it has been shown that biomass maximising flux states are usually degenerate, in that exist multiple flux distributions that yield the same maximal biomass value [13].

The limitation can be tackled using a multi-objective optimisation scheme; in this context the results of the optimisation process is a set of non-dominated points, called the Pareto front. As shown by Costanza *et al*. [4] the multi-objective optimisation problem can be tackled using bilevel linear programming coupled with evolutionary algorithms. In this work we choose the following bilayer formulation, as proposed before [1]:

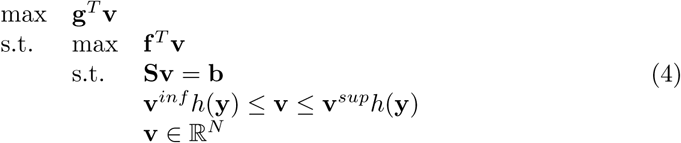

being *f* and *g* are arrays of weights associated with the first and second objectives respectively and being *h*(**y**) a multiplicative function of gene expression vector y, introduced below.

In order to exploit multiple source of information, we use a multi-omic model, integrating gene expression data into the model. Each metabolic reaction in the model depends on a set of genes through gene to protein to reaction (GPR) associations [21]. Each GPR is described as a string of genes linked by AND/OR operators; if a gene set is made up of two genes connected by an AND, both genes are necessary to carry out the reactions, thus they made up an enzymatic complex. Conversely if two genes are connected by an OR operator they independently catalyse the same reaction, making up an isozyme. In order to deal with continuous expression values, instead of binary (zero or one) activations, we borrow the standard operators from fuzzy logic [31]. The expression of two genes connected through an AND operator is the minimum of the expression of the individual genes making up the gene set. The gene set expression for two genes connected by an OR operator is the max of expression of the individual genes.

The expression of a get set can then be applied to the corresponding reaction flux boundaries using the following piece-wise multiplicative function:

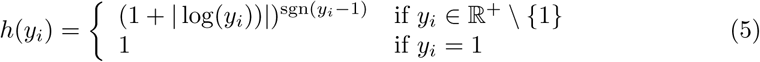

being *y_i_* the expression of the gene set responsible for the *i*-th reaction [14]. The function approximate the relationship between mRNA abundance and protein synthesis rate [8]. A discussion of the appropriateness of using mRNA levels as a proxy for protein abundance is given elsewhere [1].

### 2.2 Approximating FBA

In order to be able to train a VAE to reconstruct a GE vector that produces the same fluxes as the input one, we decided to use the mean square loss between the fluxes of the reconstructed GE, let’s say *F*′ and the fluxes of the original GE vector, *F*; nevertheless, doing so requires the FBA layer in Figure 1 to be differentiable with respect to the GE. To the best of our knowledge no differentiable formulation of linear programming w.r.t a non-differentiable function of the constraints has been proposed before. Thus, we approximated the FBA using a 5-layered MLP (APPROX. FBA in Figure 1); the architecture is detailed below.

The training is schematised in Figure 2; we provided the GE vector, as well as the concentration of relevant metabolites (glucose and oxygen) in the media as inputs. It is worth underlining the the learnt model is limited to a particular organisms and to variations of metabolite concentrations for which it has been trained; notably, the model provides a sound approximation only for GE vectors that reside in the convex hull of the training data. Nevertheless, this aspect does not represent a limitation for the current purpose of the model (See Section 3.1). The FBA approximator is trained independently from the remaining architecture, and its weights are kept fixed when training the VAE.

**Figure 2.**
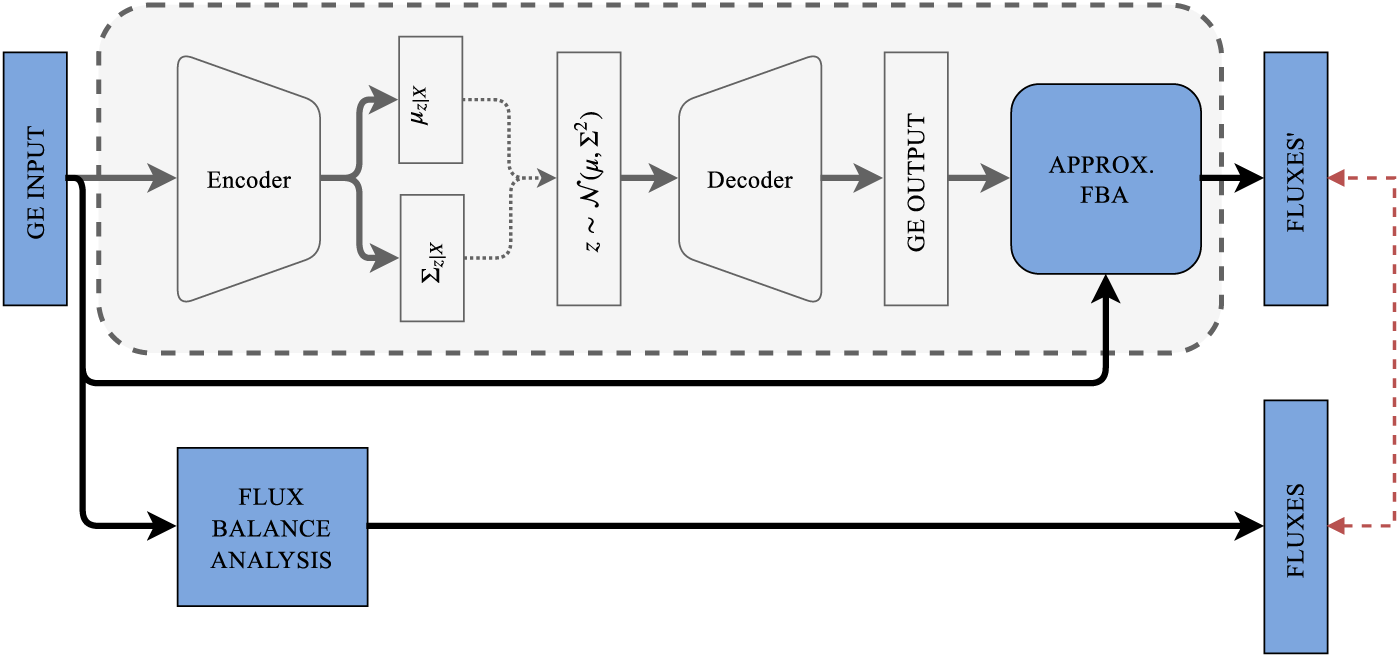
Scheme of training process for the FBA approximator. We provide a GE vector, as well as the concentration of relevant metabolites in the media (oxygen and glucose) as inputs to the FBA approximator; the approximator is trained to approximated the fluxes obtained via FBA.

The FBA approximator is 5-layered MPL; its architecture is shown in Figure S1. The total number of trainable parameters is 24,337,944. We used *relu* as activation functions for all but the last layer; the last layer uses a linear activation function. We used an adagrad optimizer, and mean square error (MSE) between the reconstructed and FBA-produced layer as loss function. We trained the network on a NVIDIA TESLA K80 GPU; we trained for 300 epochs, using a batch size of 128.

### 2.3 Dataset

#### 2.3.1 Real Dataset

As a real dataset we use a compendium of 466 *E. coli* Affymetrix Antisense2 microarray expression profiles [6]. The dataset includes gene expression microarrays collected in different experimental conditions, such as varying oxygen and glucose concentrations, pH changes, heat shock and antibiotics. When running the FBA, we consider oxygen and glucose intake rate depending on the particular media condition.

#### 2.3.2 Generating GE data

In order to construct a dataset we used a multi-objective optimisation algorithm, generating a set gene expression vectors defining Pareto fronts for different experimental conditions (aerobic/anaerobic, varying concentration of glucose in the medium). We generated, for a set of 12 experimental conditions, Pareto fronts using the METRADE multi-objective scheme [1], with 450 iterations and a population of 300 individuals. METRADE couples a bilayer FBA formulation with a NSGA-II algorithm [5]. We followed a previously proposed approach [14], selecting biomass maximisation and total intracellular flux minimisation as conflicting objectives.

We selected various experimental conditions: anaerobic and aerobic, for different glucose concentrations (5.5, 5.6, 6 10, 11, 20, 22 and 44 mmol h^−1^ gDW^−1^), in order to provide a sufficient number of Pareto fronts. The experimental conditions span the set of glucose and oxygen concentrations found in the dataset of 466 *E. coli* Affymetrix Antisense2 microarray expression profiles described before. The objective space for a sub-sample of the dataset is shown in Figure S4. The resulting dataset is composed of 6’440 gene expression vectors.

### 2.4 VAEs

Variational autoencoders (VAEs) [17] are a class of deep generative models trained via variational inference methods, hence the name. Given a dataset 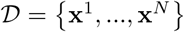 a probabilistic encoder and decoder are trained. The generative process considers a set of latent variables **z** ∈ ℝ*^k^* and a set of observed variables 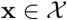, thus defining a joint probability distribution *p*(**x**, **z**) = *p*(**x**|**z**)*p*(**z**).

We want to maximise the marginal probability of each x in the training set under the generative process 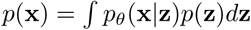; nevertheless, being the integral intractable, VAEs are generally trained by maximising the evidence lower bound (ELBO) instead:

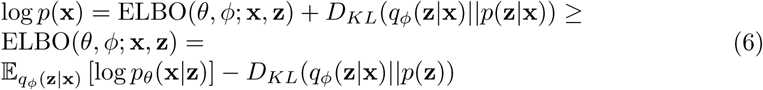

where *q_ϕ_*(**z**|**x**) is an approximate (posterior) distribution, parametrised via *ϕ*. Here 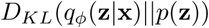 is the Kullback Leiber divergence between *q_ϕ_* and *p*.

The optimisation is achieved by simultaneously performing gradient descent on both *ϕ* and *θ*; to do so the *reparametrization trick* is used, i.e. each random variable 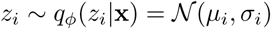 is rewritten as a differentiable transformation of a noise variable 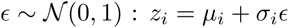. In the VAE model we generally assume 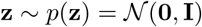 and 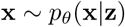; furthermore both p and q are parametrised by *neural networks*.

In this work we use a particular evolution of the VAE framework, that modify the original objective function introducing an adjustable hyperparameter *β* [11], hence the name.

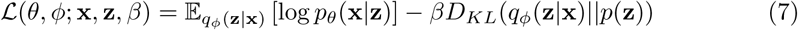

The parameter *β* determines the amount of pressure applied to the latent information channel. Imposing *β* > 1 we strongly constraints the latent bottleneck, thus pushing *q_ϕ_*(**z**|**x**) toward the isotropic unit Gaussian prior *p*(**z**). It has been shown that reconstruction under this bottleneck helps producing a disentangled representation [3], in which different latent units are sensible to changes in a single generative factor and approximately invariant with respect to changes of the other factors.

Here we briefly justify the choice of the model, providing the rationale behind the choice of the VAE instead of a more common AE; although the VAE has an AE like structure, i.e. it is made of a decoder and an encoder network, it serves a much larger purpose; still it can be used fruitfully to learn latent representations. Since we are interested in having a probabilistic generative model of the data, a VAE is a better choice. Amongst the advantages offered by a generative model, it is that of sampling new elements from the distribution, e.g. sampling fake gene expression data for a particular condition.

The overall VAE model is made of an encoder *q* and a decoder *p*. The decoder, *p_θ_*(**x**|**z**), is a 3-layered MLP as shown in Figure S2; the input to the model is the sampled latent state **z**. The encoder *q_ϕ_*(**z**|**x**) is a 3-layered MLP as shown in Figure S3. We train the VAE network on a NVIDIA TESLA K80 GPU for 300 epochs, using a batch size of 128, and keeping the approximated FBA weights fixed.

## 3 Experimental Results

In this section, we fit our framework on the dataset we have generated, evaluating its ability to learn a compact and disentangled representation of the gene expression data. We first characterise the FBA approximator (Section 3.1); later on we evaluate the GEESE model (Section 3.2). Then we compare GEESE with a baseline VAE on a real dataset (Section 3.2).

### 3.1 FBA approximation

We first trained the FBA approximator. Then, we evaluated the approximator on both the train and test set; we show the results in Figure 3. The MSEs are 0.671 and 1.048 on the train and test set respectively.

**Figure 3.**
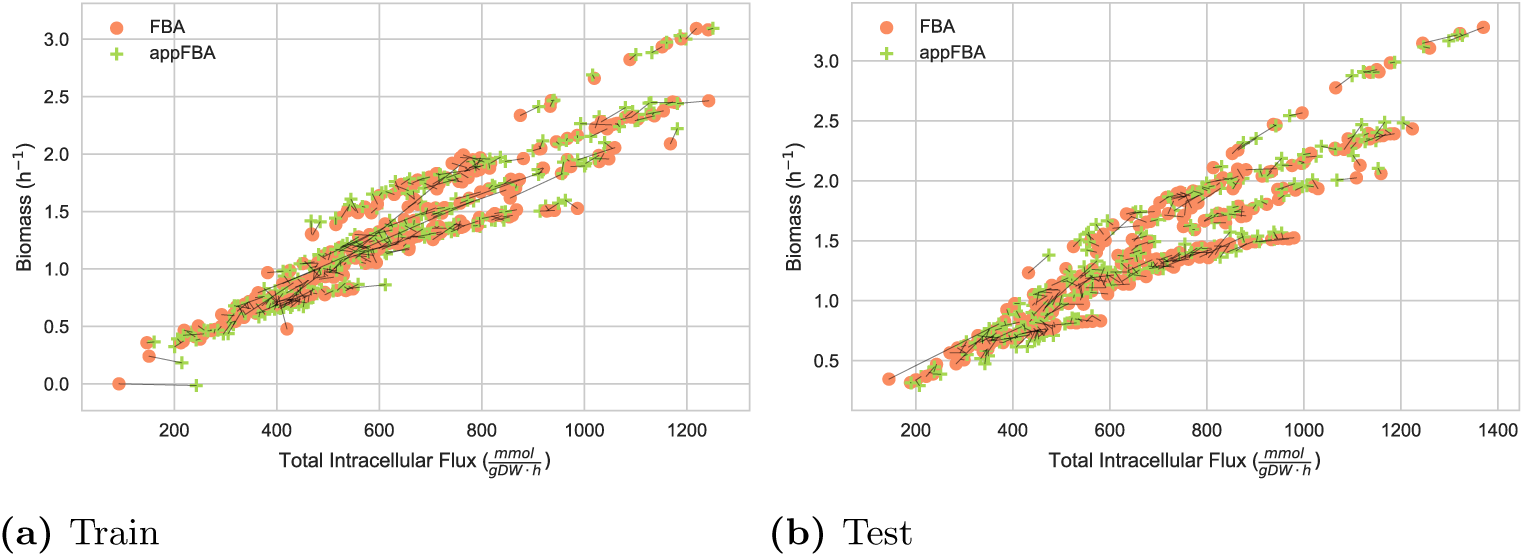
Pareto preserving properties of the FBA approximator; for sake of clarity we show only 300 random samples from both the train (upper panel) and test (lower panel) sets. The FBA predictions are represented as orange cirles, while the approximated FBA ones are marked with green + symbols. Each pair of prediction is connected by a black line.

In Figure 3 we plotted the two objectives for both FBA-generated fluxes (orange circles) and approximated FBA (green + symbols); corresponding points are connected by a black line. According to the experimental results, the approximator learn a property we dubbed *Pareto-preservation*, i.e. it generally learns to map g.e. data in the nearest (true) Pareto Front.

Furthermore, in order to asses the deviation w.r.t the steady-state assumption **Sv = 0**, we measured 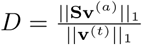, being **v**^(^*^a^*^)^ and **v**^(^*^t^*^)^ the approximated and exact flux vectors respectively; we will refer to *D* as the deviation index hereafter. The mean deviation indexes on the train and test set are 0.119 and 0.115 respectively.

It is worth noticing that, while the FBA approximator shows promising results, we are not interested in obtaining a perfect approximation of the FBA; as long as the approximator drives the latent space learning in the direction of a better feature extraction, the approximation is good enough. Whether or not this property is full-filled is the subject of the following subsections.

Furthermore, we expect the model to generalise as long as the data on which it is used, lie in the convex hull of the training distribution [10]; since the FBA approximator is used to train the VAE only we can be reasonably confident on its behaviour.

### 3.2 Evaluating the VAE

We trained the variational autoencoder shown in Figure 1. Details about the training process and the VAE architecture are reported in Section 2.4. We then evaluated the learnt FBA on a hold out test set.

We first inspected the output gene expression reconstructed by the autoencoder; interestingly, the reconstructed gene expression data shows greater compactness: as shown in Figure S6, the standard deviation of the reconstructed gene expression (lower panel) is, for the majority of genes, lower than the original one (upper panel). This behaviour suggests that the VAE is denoising the data, discarding all the variation in gene expression that is not strictly useful in term of a reconstructing a particular metabolic phenotype.

We then evaluated the learnt latent space in terms of the original experimental conditions (See Section 2.3.2). In the left panel Figure 4 we show a scatter-plot of the encoded latent dimensions for a sample of *n* = 2000 gene expression vectors. As expected,the encoding maps different experimental conditions along different curves in the latent space. The closeness between curves reflects similarities/dissimilarities between experimental conditions. For example, the experimental conditions (Anaerobic,-20) and (Anaerobic, −22) are encoded on partially overlapping curves as are their Pareto fronts in Figure S4. More interesting, each curve is directed, in the sense that moving across a specific curve we span the corresponding Pareto front (See below and left panel of Figure 4 for further details).

**Figure 4.**
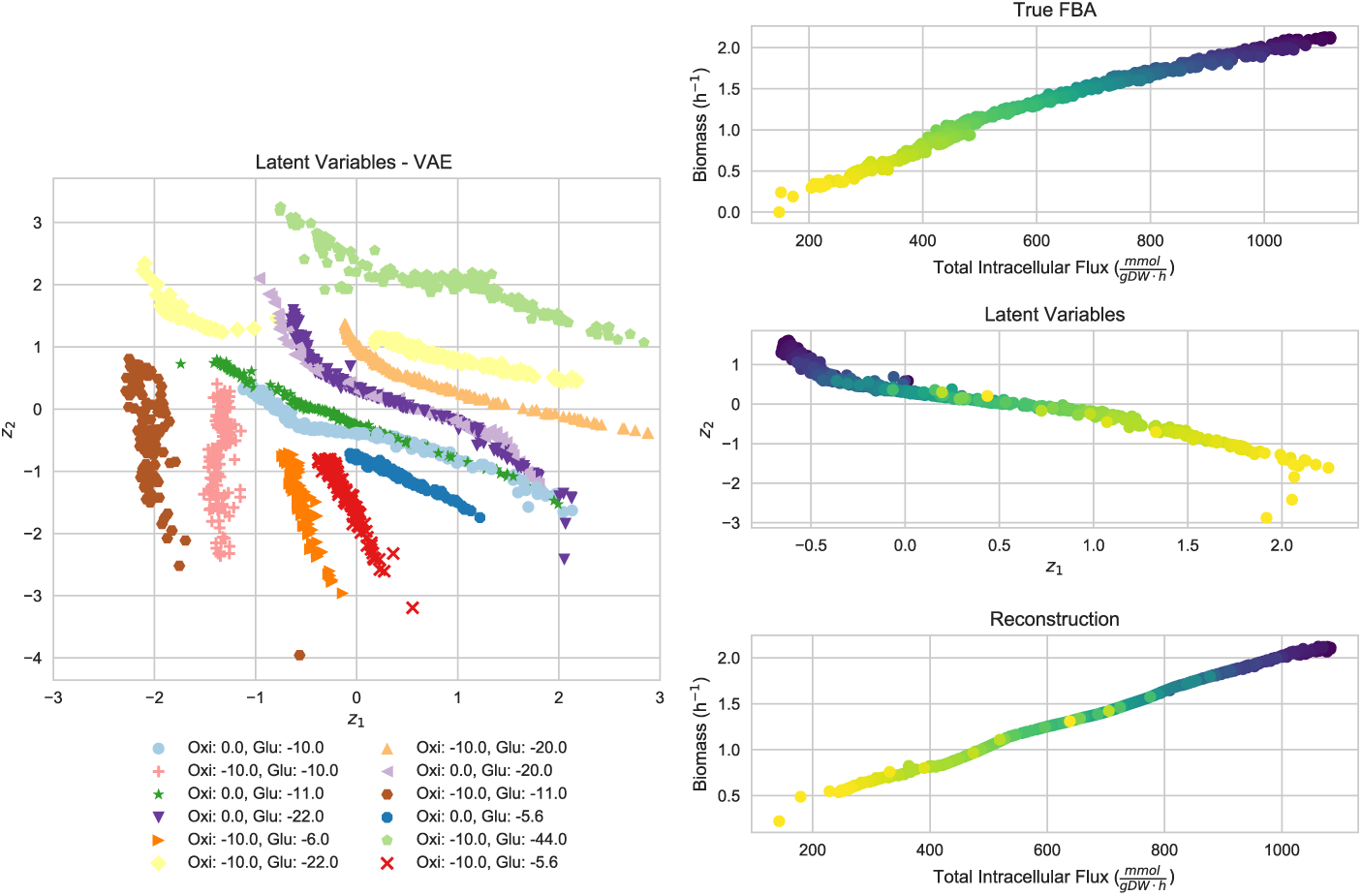
Left Panel) The (bi-dimensional) latent space for a sample of *n* = 2000 GE data is reported. The VAE encodes different experimental conditions in different regions (more properly paths) of the latent space. See also Figure S4 in supplementary material. Right Panel) For a given Pareto Front (Anaerobic, max glucose intake 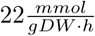), dark purple in Figure 4) we plot the two objectives for the true FBA (upper right panel) and the two objectives of the reconstructed FBA, obtained passing the gene expression data through the autoencoder (lower right panel). The colour, shifting from yellow to dark purple, encodes the the movement along the Pareto front (middle right panel).

We then compared the latent space generated by GEESE with the one induced by a baseline VAE (without the FBA module); even if the experimental conditions are nicely separated in the latent space, the Pareto front structure is not preserved (Figure S5). Indeed, even if the Pareto front structure is an inherent property of the dataset, it is only captured adding the metabolic constraints imposed by GEESE.

Furthermore we analysed the capability of the proposed framework w.r.t. the reconstruction of a given Pareto front. We selected gene expression data from a particular Pareto front (obtained in anaerobic setting with max glucose intake 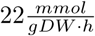)), and we used our VAE, trained with a latent space dimension of 2. In the right panel of Figure 4 we report the objective space produced by the true FBA (upper right panel), the encoding of the gene expression data into the latent space (dark purple curve in the left panel), and the two objectives of the reconstructed FBA, obtained passing the gene expression data through the autoencoder (lower left panel).

We analysed the capability of the proposed framework by locally exploring the adjacent latent space of the given Pareto fronts. New data points were generated in the proximity of a particular Pareto front (obtained in anaerobic setting with max glucose intake 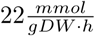)), followed by the reconstruction of their Gene Expression using our VAE (left panel in Figure 5). The distribution of each gene was used to identify a set of particularly stable and unstable genes along that region of the latent space. We performed sensitivity analysis to obtain the robustness value of each gene for two different objectives. The genes classified as stable in the local latent space around the Pareto front show high values of robustness, meaning that their role on the optimisation under those specific experimental conditions is replaceable (See right panel of Figure 5). Instead, the unstable genes show mixed robustness values, which can be understood as having a more critical influence over the optimisation under these experimental conditions. In both groups, the VAE manages to identify genes that have almost binary optimisation values (either completely breaking or maximising the results).

**Figure 5.**
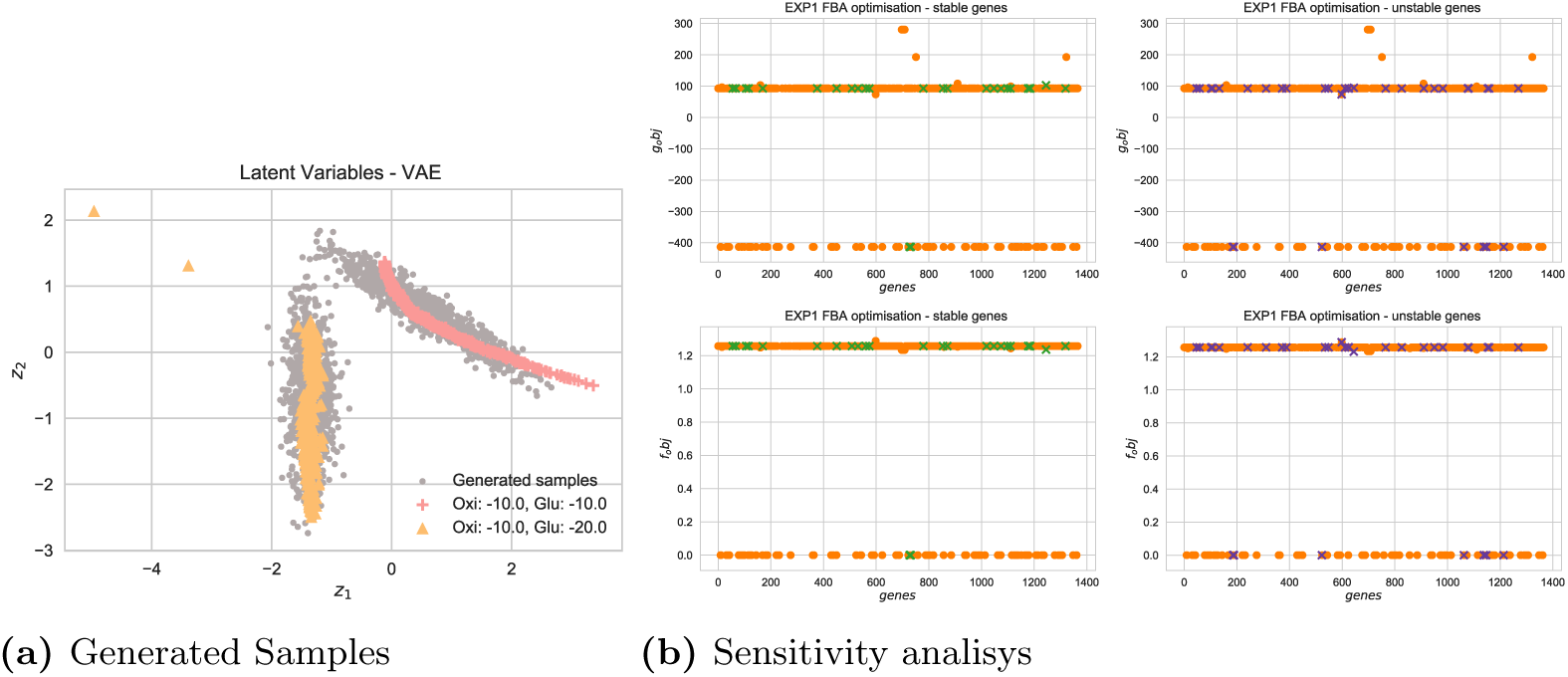
New data points were generated in the proximity of a particular Pareto front (left panel) followed by the reconstruction of their Gene Expression using our VAE. Stable and unstable genes have been identified, and sensitivity analysis for each gene for two different objectives, have been performed (right panel). In both groups, the VAE manage to identify genes that have almost binary optimisation values.

Therefore VAE succeed in identifying a set of specific genes that are characteristic to each of the experimental conditions, with minimal overlap among pareto fronts and clear effect over optimisation.

### 3.3 Is GEESE helping in reconstructing the latent space?

In order to inspect how the model generalise in a real case scenario, we devised the following experiment: we selected a compendium of 466 *E. coli* Affymetrix Antisense2 microarray expression profiles [6]. As described below, this dataset collects gene expression microarrays from different experimental conditions, such as varying oxygen and glucose concentrations, pH changes, heat shock and antibiotics. We then generated a dataset, using only the knowledge of glucose and oxygen in the media, using Paretos from the multi-objective evolutionary approach. We trained both a baseline Variational Autoencoder, without the FBA module, and our architecture (GEESE), on this dataset. Both GEESE (Figure 4) and the baseline VAE (Figure S5), show nice properties in the embedded space.

Then, considering only the learnt encoder as a non-linear feature extractor, we inspected the latent space for the selected (unseen) compendia of real gene expression data. The embeddings are reported in Figure 6a and in Figure 6b for the VAE and GEESE respectively.

**Figure 6.**
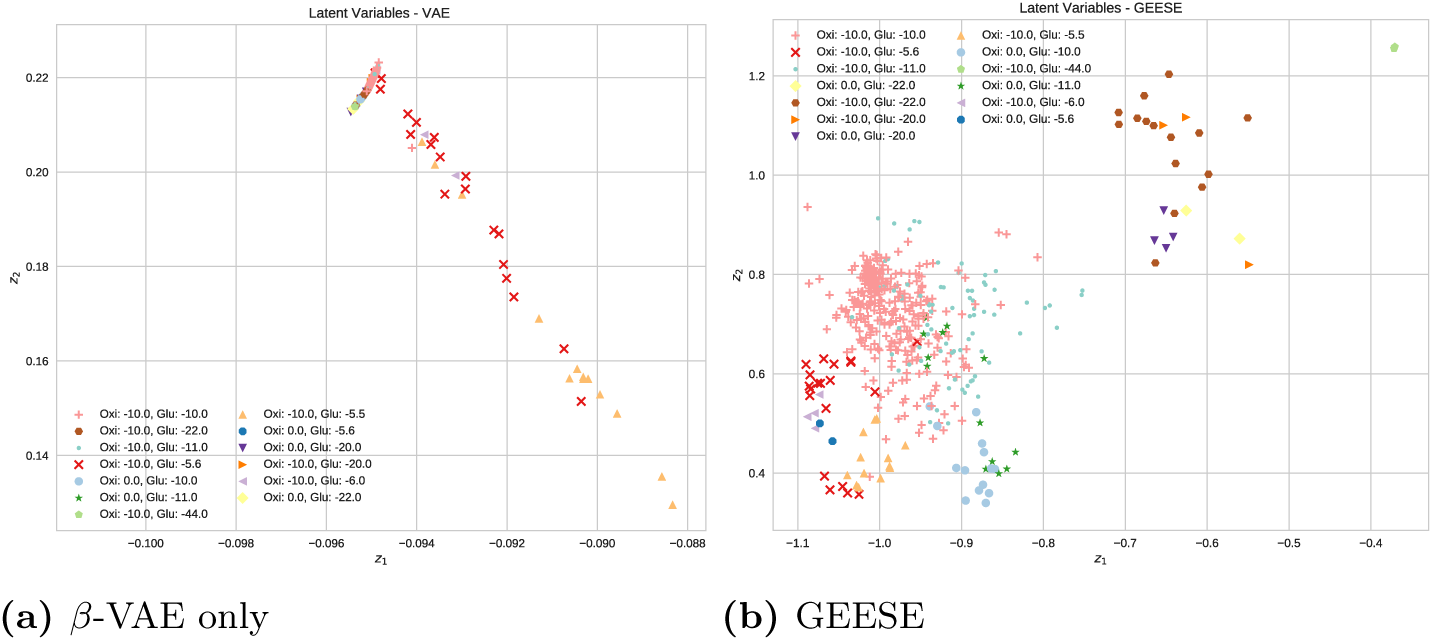
The latent spaces for (unseen) examples from the real Dataset are shown for a baseline VAE and our proposed architecture (GEESE).

Interestingly, the GEESE architecture shows a greater capability to generalise to unseen new data, and it is able to better spread the unseen data in the latent space. The baseline VAE, instead, collapse the majority the new data in very thin region of the latent space. We hypothesise that, in this context, the GSMM works as a prior, thus providing sounded built-in assumptions about the structure of the underlying data, and helps in make sense of noisy, high dimensional measures.

## 4 Discussion and Conclusion

A fundamental question in biology is to understand what makes individuals, populations, and species different from each other. The notion of phenotype i.e. observable attributes of an individual, is opposed to that of genotype, the inherited material encoded in genes. The ongoing quest of system biology is that of bridging the genotype-phenotype gap. Both gene expression microarrays and RNA-seq have been used to probe the transcriptional landscape of a biological sample. Notwithstanding, the prediction of a metabolic phenotype from gene expression levels is anything but a solved problem.

In this work we tackled the problem of learning a meaningful latent space for gene expression data, using a deep generative model. In order to force the latent space to convey metabolic information, we devised a specific framework, composed of a generative deep neural network and an approximation of a metabolic network based flux balance analysis. We first trained the FBA approximator as a 5-layered MLP; we then used a *β*-Variational Autoencoder (*β*-VAE), and we trained the model to reconstruct a gene expression vector that passing through a FBA produces the same fluxes distribution as the input one.

The learnt bi-dimensional manifold provides meaningful insights into the relationships between samples; first, the autoencoder is able to recognise gene expression vectors associated with different experimental conditions (e.g. aerobic/anaerobic and different values of glucose in the medium). Second, the autoencoder denoises the input data, while preserving the same metabolic phenotype. Third, the encoder map a Pareto front of gene expression vectors into an oriented path in the bi-dimensional space (Figure 4). The oriented paths depict how the bacteria benefits from using different GE strategies to cope with varying environmental conditions; the overlapping between strategies, notably when the environmental conditions are similar, buttresses the biological soundness of the inferred results. Indeed, the comparison with Figure S5 clearly shows how this information cannot be extracted by means as a standard AE. Finally, evaluating the learnt model on a unseen real dataset of gene expression microarray, we shows that the proposed model achieve a better generalisation with respect to a baseline VAE, as pictured in Figure 6.

We state that constraining the reconstructed gene expression to approximate the original FBA output, we are driving the autoencoder to incorporate prior knowledge about the correlations between genes and, thus, helping in make sense of noisy, high dimensional measures. The devised framework, dubbed GEESE (Gene Expression latEnt Space Encoder), represents a step towards the understanding of the genotype-phenotype relationship. As GSMMs are becoming available for an increasing number of organisms, the applicability of the proposed model is broadening. Finally, we believe that our model provides better understanding of different relevant conditions, and provides a novel perspective of the problem of manifold learning in general.

## Supplementary Figures

**Figure S1.**
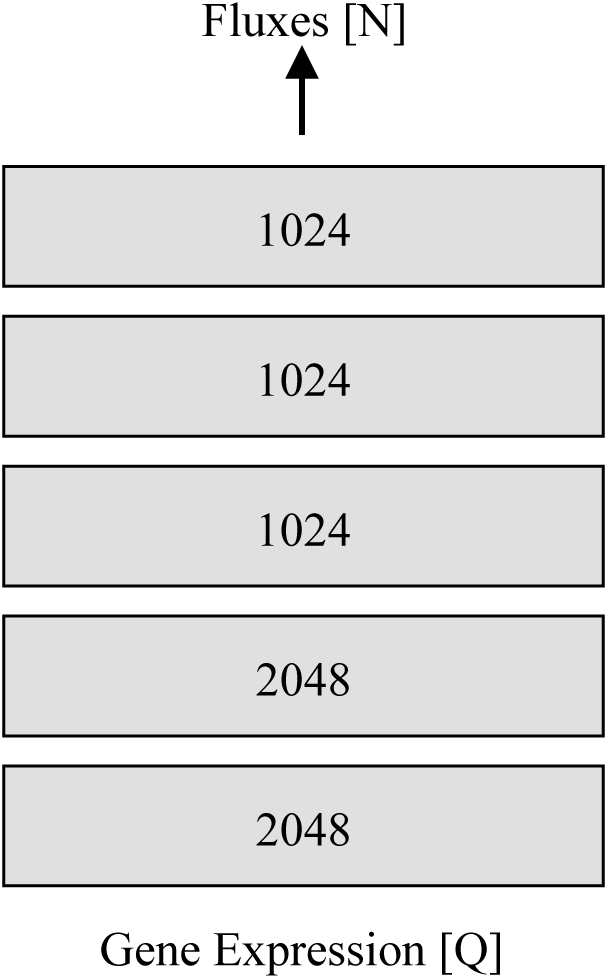
Architecture of the FBA approximator

**Figure S2.**
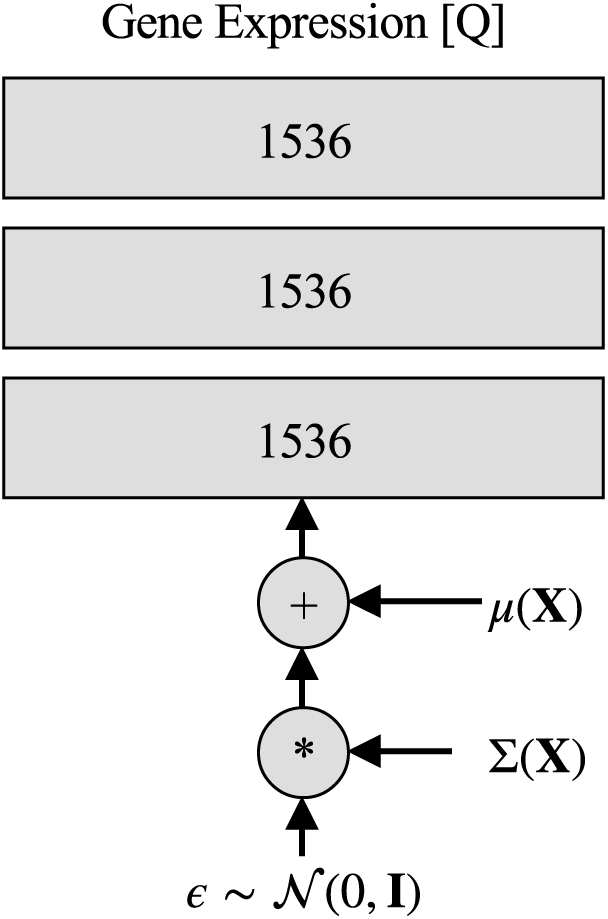
Architecture of the decoder network.

**Figure S3.**
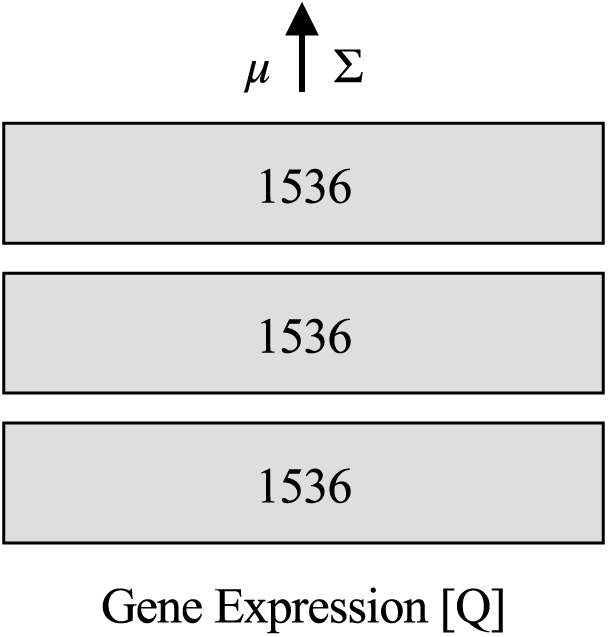
Architecture of the encoder network.

**Figure S4.**
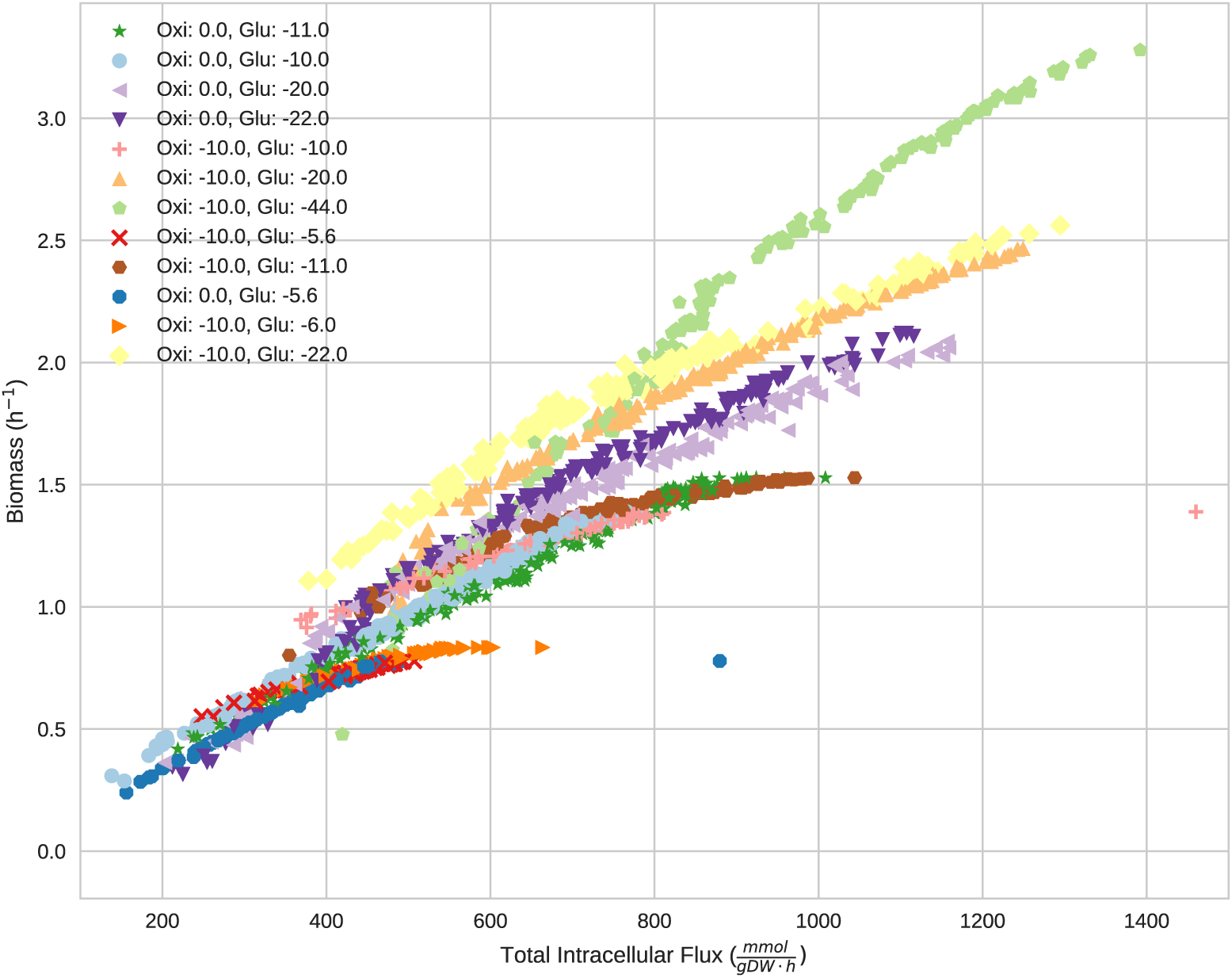
Plot of the two objectives for a sample of *n* = 2000 points from the dataset.

**Figure S5.**
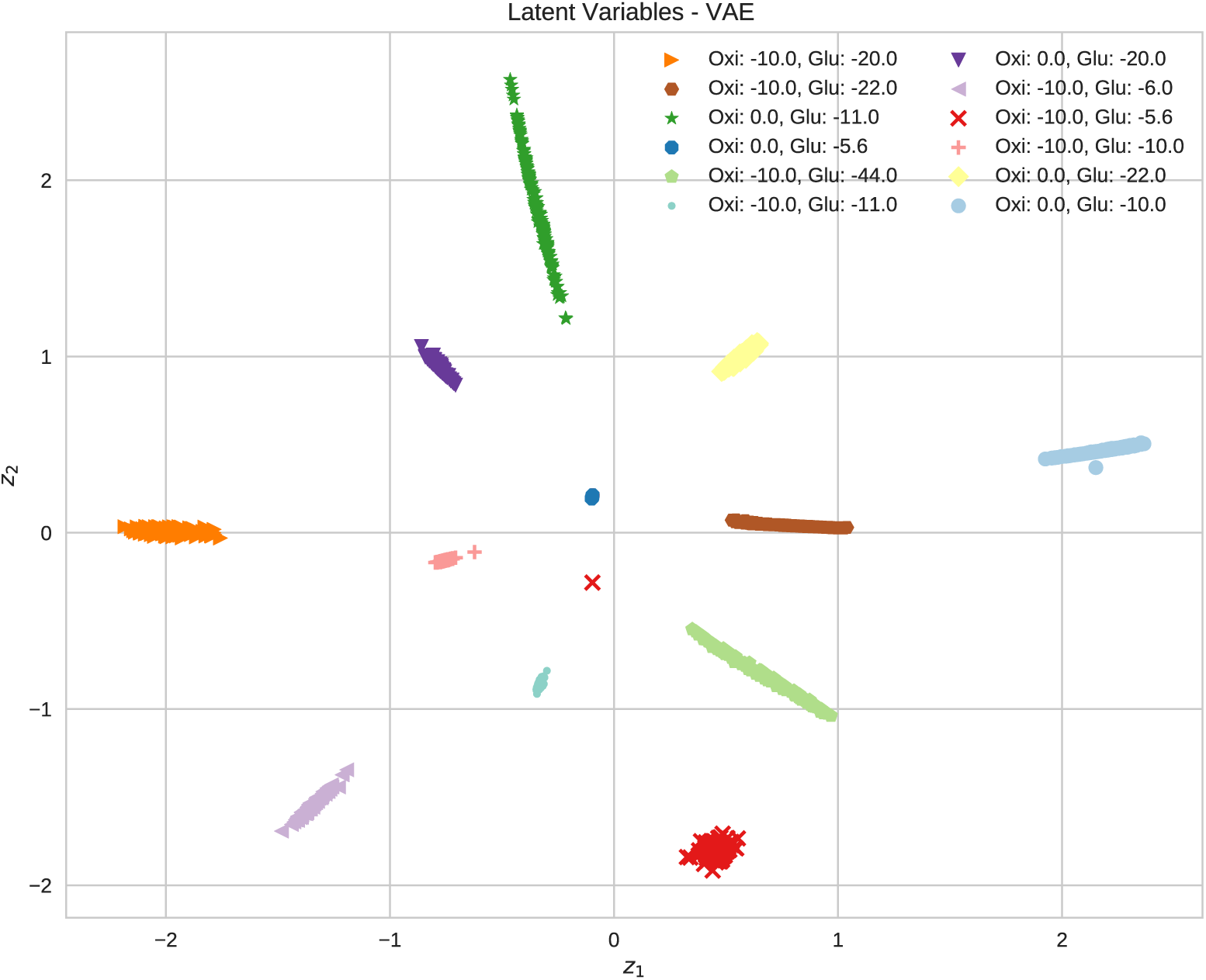
The latent space for a sample of *n* = 2000*GE* data is reported; the data have been generated using the VAE only. The VAE still encodes different experimental conditions in different regions of the latent space, but the path-like structure is missing.

**Figure S6.**
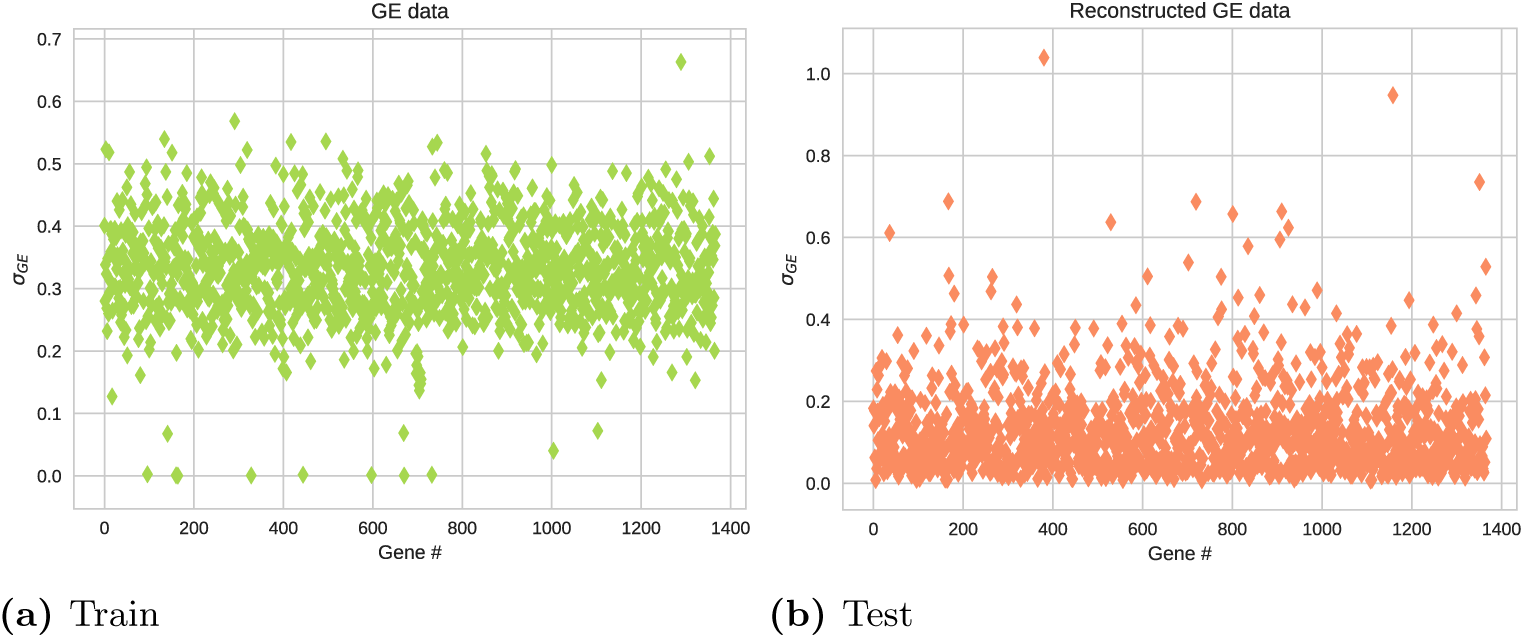
Standard deviation of both the gene expression data (upper panel) and the reconstructed gene expression data (lower panel) across the dataset. The reconstructed gene expression is more compact.

